# Stabilizing the dynamics of laboratory populations of *Drosophila melanogaster* through upper and lower limiter controls

**DOI:** 10.1101/023507

**Authors:** Sudipta Tung, Abhishek Mishra, Sutirth Dey

## Abstract

Although a large number of methods exist to control the dynamics of populations to a desired state, few of them have been empirically validated. This limits the scope of using these methods in real-life scenarios. To address this issue, we tested the efficacy of two well-known control methods in enhancing different kinds of stability in highly fluctuating, extinction-prone populations of *Drosophila melanogaster*. The Upper Limiter Control (ULC) method was able to reduce the fluctuations in population sizes as well as the extinction probability of the populations. On the negative side, it had no effect on the effective population size and required a large amount of effort. On the other hand, Lower Limiter Control (LLC) enhanced effective population size and reduced extinction probability at a relatively low amount of effort. However, its effects on population fluctuations were equivocal. We examined the population size distributions with and without the control methods, to derive biologically intuitive explanations for how these control methods work. We also show that biologically-realistic simulations, using a very general population dynamics model, are able to capture most of the trends of our data. This suggests that our results are likely to be generalizable to a wide range of scenarios.

## 1. INTRODUCTION

Over the last two decades, several methods have been suggested in control theory (Chernousko et al. 2008) and theoretical nonlinear dynamics (Andrievskii and Fradkov 2003, 2004, Schöll and Schuster 2008) to stabilize unstable non-linear dynamical systems. Several of these methods perturb system parameters in real time to attain desired behaviours like stable points or simple limit cycles (Garfinkel et al. 1992). Unfortunately, for even fairly simple biological populations, the exact equations underlying the dynamics are often unknown. Moreover, when available, the parameters of such equations (e.g. carrying capacity or intrinsic growth rate) can often only be estimated *post-facto* through model-fitting and thus are not available for real time perturbations. Finally, due to the ubiquity of noise in biological systems, it is not only impossible to attain stable points or limit cycles in the strict mathematical sense, it also becomes very difficult to distinguish such behaviours from chaotic dynamics (although see Desharnais et al. 2001). Thus, a different class of control methods and observables are needed in the context of biological populations.

The choice of method also critically depends upon the desired goal of control. There are two major, typically mutually exclusive, motivations for stabilizing biological populations. The first is in the context of economically exploited species (e.g. fishes) where the aim is to maximize the yield over a long period of time and reduce the uncertainty of the yield (Lande et al. 1997). The second aim seeks to reduce the amplitude of fluctuation in sizes or increase the long term probability of persistence for populations (Hilker and Westerhoff 2007, Gusset et al. 2009). Not surprisingly, stabilizing the yield of harvested populations has received far more theoretical and empirical attention (Milner-Gulland and Mace 1998) than stabilizing threatened species. Part of the problem with the latter is that conservation efforts are often directed towards charismatic species of mammals and birds whose dynamics are much more complicated than the simple models often studied by population dynamists. However, it should be noted that a simple model like the Ricker model (Ricker 1954) does provide fairly accurate descriptions of the dynamics of taxonomic groups including bacteria (Ponciano et al. 2005), fungi (Ives et al. 2004), ciliates (Fryxell et al. 2005), insects (Sheeba and Joshi 1998) and fishes (Denney et al. 2002). Together, such taxa account for a huge fraction of the total biodiversity on earth, at least some part of which have already been recorded to be extinct (Baillie and Butcher 2012). Therefore, there is a need to study the methods that can stabilize the dynamics of such “non-charismatic” taxa.

A major hindrance in applying the insights gained from theoretical studies in controlling endangered populations is the fact that few of the proposed methods have been empirically validated even under laboratory conditions (however see Desharnais et al. 2001, Becks et al. 2005, Dey and Joshi 2007), let alone in nature. Given that survivals of threatened species are at stake, it is understandable when practitioners of conservation are unwilling to try out untested methods in the field. On the other hand, new methods have to be validated somehow in order to assess their suitability for a given scenario. A reasonable way out of this impasse is to validate these methods under laboratory conditions. The success of a method to stabilize laboratory populations allows us to verify our understanding about how the method works. Unfortunately, it does not guarantee the method’s success under field conditions but merely increases the confidence that can be placed on its success. On the other hand, the failure of a method under laboratory conditions would typically suggest lack of understanding regarding some crucial aspect of the biology of the system.

In this study, we use unstable laboratory populations of the common fruit-fly *Drosophila melanogaster* to investigate the stabilizing properties of two control methods, namely upper limiter control (ULC) and lower limiter control (LLC). For each of these control methods, we investigate two different arbitrarily chosen values of the controlling parameter. ULC and LLC belong to the broad classes of methods that involve culling and restocking respectively. We chose these two over many such available culling / restocking schemes primarily because they have been extensively investigated theoretically and numerically (Hilker and Westerhoff 2005, 2006, Tung et al. 2014). This means that a number of predictions already exist in the literature for verifying against our empirical data. Therefore, the main focus of this paper was on an intuitive understanding of how these two methods affect the dynamics. We understand that there might be other culling / restocking methods that might have also been extensively investigated theoretically. However logistic considerations in terms of running the experiments constrained us to only two methods.

Here we show that ULC reduces temporal fluctuations in population sizes, as well as the extinction probability of populations. However, it is unable to enhance the effective population size and has high effort magnitude. On the other hand, the efficacy of LLC in reducing the fluctuations in population sizes is equivocal. In spite of that, the method is able to cause significant reduction in extinction probability and increased effective population size. Most importantly, the effort magnitude required to stabilize the populations is much less compared to ULC. We provide biologically intuitive explanations of how these control methods stabilize the populations. We also experimentally verify several theoretical predictions from the literature and show that our empirical results agree well with biologically realistic simulations.

## 2. METHODS

### 2.1. Maintenance regime of the flies

In this study, we used individuals from a large (breeding size of ∼2400) laboratory population of *Drosophila melanogaster* called DB_4_. The detailed maintenance regime and ancestry of this population has been described elsewhere (Sah et al. 2013). From this population, we derived 30 single vial cultures, each of which represented an independent population. Each of these populations was initiated by placing exactly 10 eggs on 1.1 ml of banana-jaggery medium in a 30 ml plastic vial. The vials were placed in an incubator at 25°C under constant light conditions. Once eclosion started, the freshly emerged adults of a population were daily transferred to a corresponding adult-holding vial, containing approximately 6 ml of banana-jaggery medium. This process continued till the 18^th^ day after egg-collection, after which the egg-vials were discarded. The adult flies were then supplied with excess live yeast paste for three days to boost up their fecundity. On the 21^st^ day after egg-collection, the adults were counted and culling or restocking of flies was imposed as per the prescribed control regimes (see section 2.2). Since the dynamics of a sexually reproducing species is primarily governed by the number of females, culling or restocking was implemented only on the female flies (Dey and Joshi 2006, Dey and Joshi 2007). The adults were then allowed to oviposit in a vial containing 1.1 ml of medium for 24 hrs. After oviposition, the adults were rejected and the eggs formed the next generation. The experiment was run over 14 generations. Theoretical (Mueller 1988) and empirical (Dey and Joshi 2006, Sah et al. 2013) studies have shown that a combination of low levels of larval food (1.1 ml here) and excess live yeast paste destabilizes the populations by inducing large amplitude oscillations in the time series. This nutritional regime thus allowed us to study the stabilizing effect of various control methods on populations whose dynamics were otherwise unstable.

### 2.2 Control methods

Upper Limiter Control (ULC) involves culling to a fixed threshold, i.e. the population size is not allowed to go beyond an upper value (Hilker and Westerhoff 2005). Mathematically, this is written as *N*_*t*_′ = *min (N*_*t*_, *U)* where N_t_ and N_t_′ refer to the population sizes before and after the application of the control method, *U* is the pre-determined value of the upper threshold and *min* is the minimum operator. To impose ULC experimentally, we culled the number of females in a population to the arbitrarily set levels of 15 (U1) or 10 (U2). When the number of females in a population was less than the threshold, the population was left unperturbed. Note that for ULC, lower values of U represent more stringent control and therefore U2 is a stronger control than U1.

Lower Limiter Control (LLC) is achieved by restocking the population to a fixed number, i.e. the population size is never allowed to fall below a fixed limit (*L*). Mathematically, this is given as *N*_*t*_*′* = *max (N*_*t*_, *L)* where *L* stands for the fixed lower threshold and *max* is the maximum operator. For experimental implementation, we chose two arbitrary lower thresholds of 4 (L1) and 10 (L2) females where L2 represents a stronger control than L1. Following an earlier protocol (Dey and Joshi 2006), the flies were counted, the number was multiplied by half (i.e. assuming equal sex ratio) and rounded up to estimate the number of females in the population. If this number was greater than the pre-determined value of L (i.e. 4 or 10) then the population was left untouched, else the shortfall was made up by adding the required number of females from outside. Thus, we explicitly incorporated some degree of noise in terms of application of LLC (see section 4.2 for the rationale of the same).

We used 5 replicate single vial cultures each for U1, U2, L1 and L2. We also had two batches of unperturbed populations (designated C1 and C2), each consisting of 5 replicate populations. C1 served as the unperturbed experimental controls for the ULC treatments while C2 were the corresponding experimental controls for the LLC treatments. The pairing of C1 with ULC and C2 with LLC was done *a priori*, i.e. at the time of initiation of experiments, before the data analysis. An extra set of experimental controls ensures that the ULC and LLC experiments are completely independent of each other. It also allowed us to verify the reproducibility of the unperturbed dynamics.

In case of U1, U2, C1 and C2, an extinct population was rescued by adding 4 males and 4 female flies from outside (Dey and Joshi 2006). No such intervention was needed for the L1 and L2 populations where the control method automatically ensured that the breeding population never went extinct. The flies used for rescue were maintained similarly as the unperturbed experimental populations, except that they were raised on ∼5ml of larval medium instead of 1.1 ml.

### 2.3. Constancy and persistence stability

Populations with larger fluctuations over time are deemed to have less ‘constancy’ stability and vice versa (Grimm and Wissel 1997). We used the Fluctuation Index or FI (Dey and Joshi 2006) to quantify the constancy stability of a given time series.

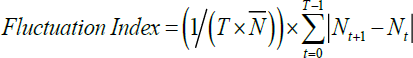

where *N*_*t*_ is the breeding population size, 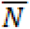 represents average population size and T denotes the length of the time series.

Persistence stability (Grimm and Wissel 1997) of a population was scored as the extinction probability during the course of the experiment (= number of extinctions / length of the time series). Since one of the methods (LLC) involved restocking, persistence was scored using the pre-perturbation census size.

### 2.4. Average population size and effective population size

Average population size 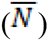 was simply the arithmetic mean of the time series while effective population size (*N*_*e*_) was quantified as the harmonic mean of the time series (Allendorf and Luikart 2007).

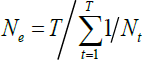

### 2.5. Effort magnitude

Following a previous study (Hilker and Westerhoff 2005), we assumed that the number of organisms to be removed or added for implementing a control method (i.e. the so called effort magnitude) is a proxy for the corresponding economic cost of implementation. Thus,

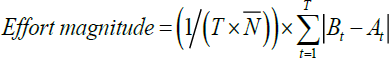

where *B*_*t*_ and *A*_*t*_ represent the pre- and post-control population sizes respectively in the *t*^*th*^ generation. 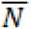 and *T* denote the average population size and length of the time series respectively (Sah et al. 2013). It should be noted here that effort magnitude is the actual number of individuals culled from or added to the population. Although the value of the *a priori* set threshold does have a bearing on the empirically observed effort magnitudes, the two are not the same quantity. Since the flies used for rescuing the extinct populations were counted as a part of effort, C1 and C2 have non-zero effort magnitudes.

### 2.6. Statistical analyses

The data for ULC and LLC were analysed separately along with their matched unperturbed populations, i.e. C1 and C2 respectively. The fluctuation index, average population size, effective population size and effort magnitude data were subjected to separate one-way ANOVA (unperturbed population and the two levels of fixed thresholds being the fixed factors). In those cases, where a significant main effect was obtained, Tukey’s HSD was employed for testing the significance of pair-wise differences among means. All ANOVA and associated comparisons were performed using STATISTICA^®^ v5 (StatSoft. Inc., Tulsa, Oklahoma). We also used the freeware Effect Size Generator (Devilly 2004) to compute Cohen’s *d* statistic (Cohen 1988) as a measure of effect sizes for the pair-wise differences among the means. The effect sizes were interpreted as small, medium or large for *d* < 0.5, 0.5 < *d* < 0.8, and *d* > 0.8.

### 2.7 Simulations

The details of the simulations are provided in section S1. In brief, we modelled each population using the Ricker model (Ricker 1954). As stated already, this model has been extensively used to represent the dynamics of wide range of organisms. The simulation framework was similar to a recent theoretical study on comparison of control methods (Tung et al. 2014).

## 3. RESULTS

### 3.1 Upper Limiter Control (ULC)

In the absence of any perturbation, the distribution of the population sizes was found to be positively skewed with a large difference between the mean and the median (Fig 1A). However, in the presence of upper thresholds beyond which the population sizes were not allowed to venture, the distributions became more symmetric, as evidenced by the low values of skew and little difference between the means and the medians (Fig 1B and 1C). This change in population distribution translated into a significant reduction in FI (F_2, 12_ = 20.8, p = 0.0001). Post-hoc tests revealed that although U1 and U2 had similar FI, both were significantly more stable than the corresponding unperturbed populations with high effect sizes (Fig 2A, Table S2). This is consistent with the fact that the population size distributions of U1 and U2 were almost identical to each other and greatly different from the unperturbed treatment (Fig 1). Interestingly, although ULC involves culling above a threshold, the effective population sizes (F_2, 12_ = 2.32, p = 0.14, Fig 2B) and the average population sizes (F_2, 12_ = 1.08, p = 0.37, Fig 2C) of the unperturbed and the controlled populations did not differ significantly (see section 4.1 for discussion). In terms of effort magnitude (Fig 2D), there was a significant main effect of treatment (F_2, 12_ = 44.07, p = 0.000003). Tukey’s HSD suggested that both U1 and U2 required significantly greater effort than the unperturbed populations and the corresponding effect sizes were large (Table S2). This is intuitive and confirms previous theoretical observations (Tung et al. 2014) that ULC requires a large amount of effort in terms of number of organisms to be removed from the controlled population. ULC also had an almost significant (F_2, 12_ = 3.61, p = 0.06) effect in terms of reduction of extinction probability (Fig 3A). The effect size of the reduction achieved by U1 and U2, compared to the unperturbed population, were found to be large (Table S2). Finally, comparing Fig 1 and 2 with the corresponding simulations (Fig S4 and Fig S5) suggests that our simulations were able to capture the broad trends of the data remarkably well.

**Figure 1.**
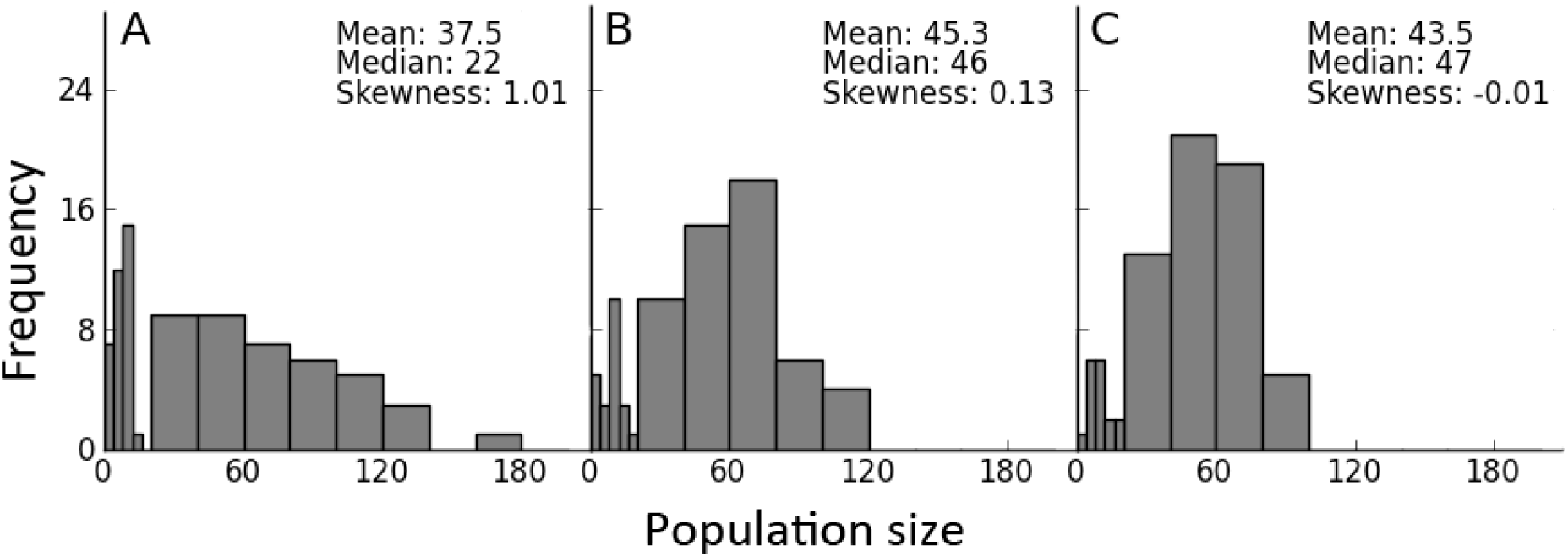
Population size distributions for unperturbed and ULC-treated populations. A. Unperturbed (C1). **B.** U1 (upper threshold = 15). **C.** U2 (upper threshold = 10). The bin size is 4 when the population size is in the range 0-20, and 20 otherwise. When the value of the upper threshold is lowered, thereby increasing control intensity, the frequency of the extreme values decreases and the overall distribution becomes more symmetric.

**Figure 2.**
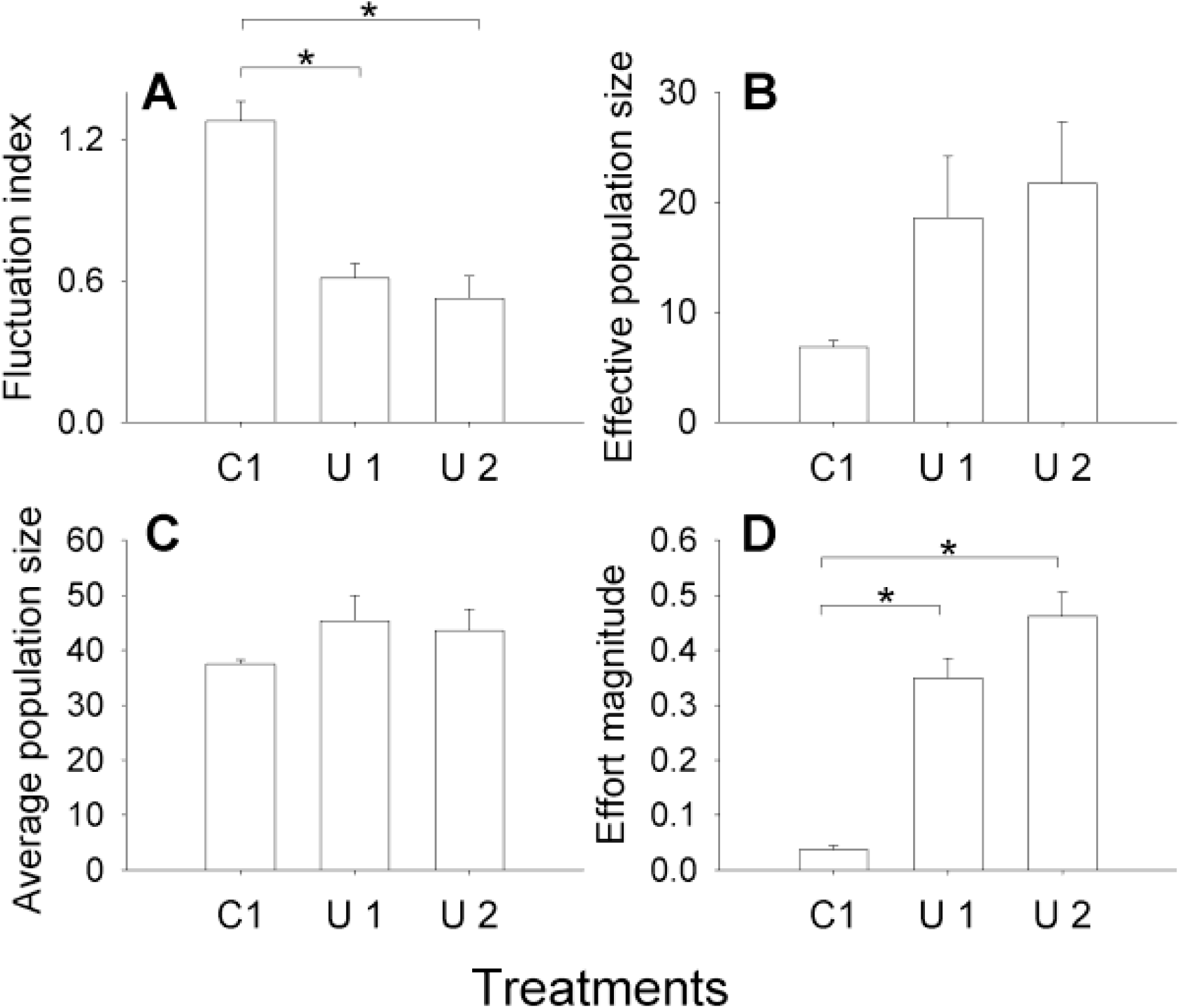
Dynamics after applying ULC. C1 represents the unperturbed populations while U1 and U2 stand for ULC thresholds of 15 and 10 respectively. Each bar represents a mean over 5 replicate populations. Error bars denote standard error around the mean and * denotes *p* < 0.05. **A. Fluctuation index**: Both the ULC treatments reduced population fluctuations significantly. Neither U1 nor U2 had a significant effect on **B**. **Effective population size** and **C**. **Average population size. D. Effort magnitude**: Both U1 and U2 required significantly more external perturbation than C1. See text for possible explanations.

**Figure 3.**
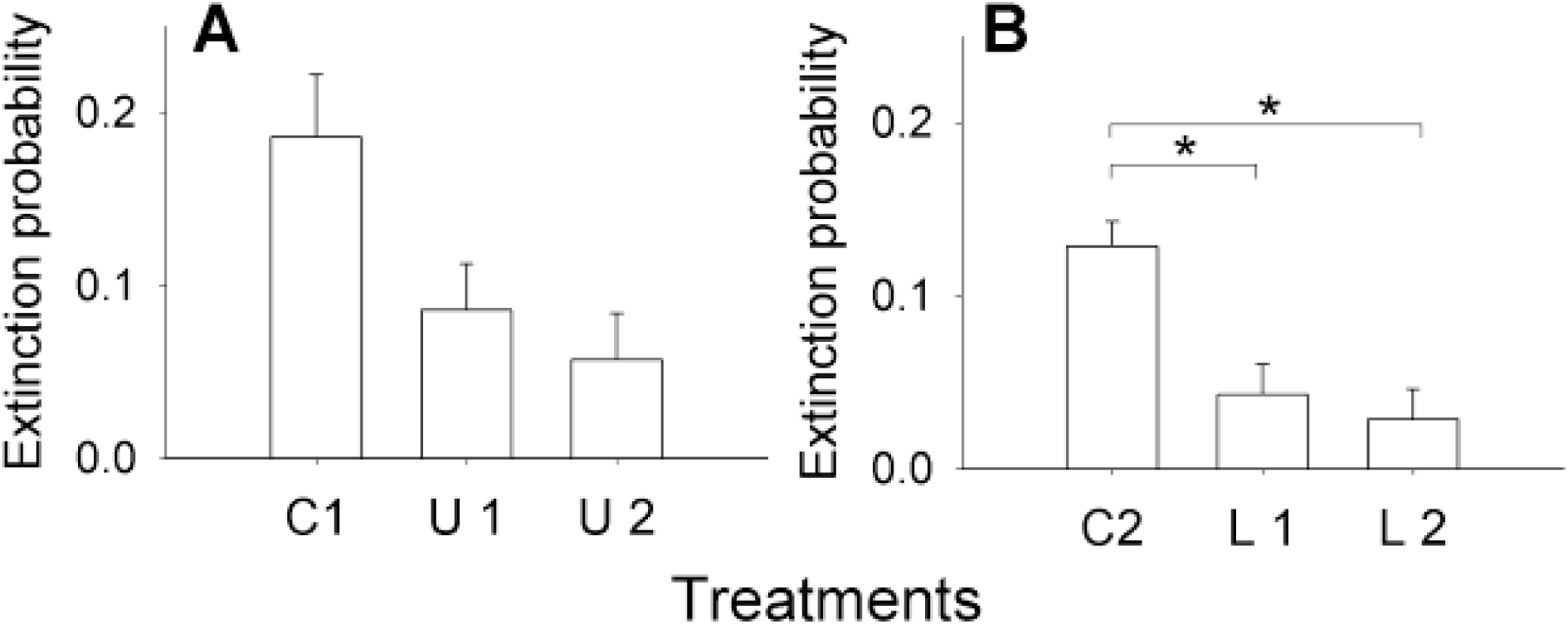
Extinction probability after applying ULC and LLC. Each bar represents a mean over 5 replicate populations. Error bars denote standard error around the mean and * denotes *p* < 0.05. **A. ULC:** C1, U1 and U2 represent the unperturbed populations, and ULC thresholds of 15 and 10 respectively. **B. LLC:** C2, L1 and L2 stand for unperturbed populations, and LLC thresholds of 4 and 10 respectively. Both control methods were able to reduce the extinction probability, although the reduction was marginally insignificant for ULC (see text for possible explanation).

### 3.2 Lower Limiter Control (LLC)

A different set of five unperturbed populations (Fig 4A) gave almost the same distributional properties as seen in Fig 1A, thus attesting the reproducibility of the distribution for the unperturbed case. However, when the population size was not allowed to fall below a fixed threshold, the changes in the population size distributions (Fig 4B and 4C) were less dramatic than those seen for ULC (Fig 1B and Fig 1C). The nature of the intervention assured that there were no individuals in the lowermost bins for the two levels of LLC (Figs 4B and 4C). Although the skew values were somewhat reduced due to the perturbations, the distribution still had a sufficiently long tail indicating that the populations were capable of reaching very high sizes. As a result of this, although there was an overall significant effect of treatment in the ANOVA (F_2, 12_ =8.75, p = 0.005), neither L1 nor L2 had significantly lower FI compared to the unperturbed treatment (Fig 5A). Interestingly though, L1 had a significantly higher FI than L2 (Fig 5A, Table S3), which is due to the fact that small values of LLC can actually increase the FI of populations (Tung et al. 2014). The effective population size of L2 was significantly higher (F_2, 12_ = 39.86, p = 0.000005) than that of the unperturbed and L1 with high effect sizes (Fig 5B, Table S2). This is intuitive as the effective population size is primarily affected by low values in a series and LLC ensures that the population size does not go below a lower threshold. However, in spite of adding individuals from outside, there was no significant difference (F_2, 12_ = 2.8, p = 0.1) between the three treatments in terms of average population size (Fig 5C). There was a significant effect in terms of effort magnitude (F_2, 12_ = 7.32, p= 0.008) and the L2 treatment required significantly greater amount of effort (with large effect sizes of the difference) compared to both L1 and the unperturbed populations (Fig 5D, Table S3). Overall, the empirical results suggest that the L1 level of LLC treatments were primarily ineffective in inducing constancy stability in the populations, while L2 was somewhat more effective (see discussion for possible explanations). However, in terms of persistence, LLC led to significant (F_2, 12_ = 5.85, p = 0.02) reduction in extinction probability. The effect size of the reduction achieved by L1 and L2, compared to the unperturbed population, were found to be large (Table S3). Finally, as with ULC, our simulations were again able to reflect the broad trends of the empirical data (*cf* Figs 4 and 5 with Figs S5 and S6 respectively).

**Figure 4.**
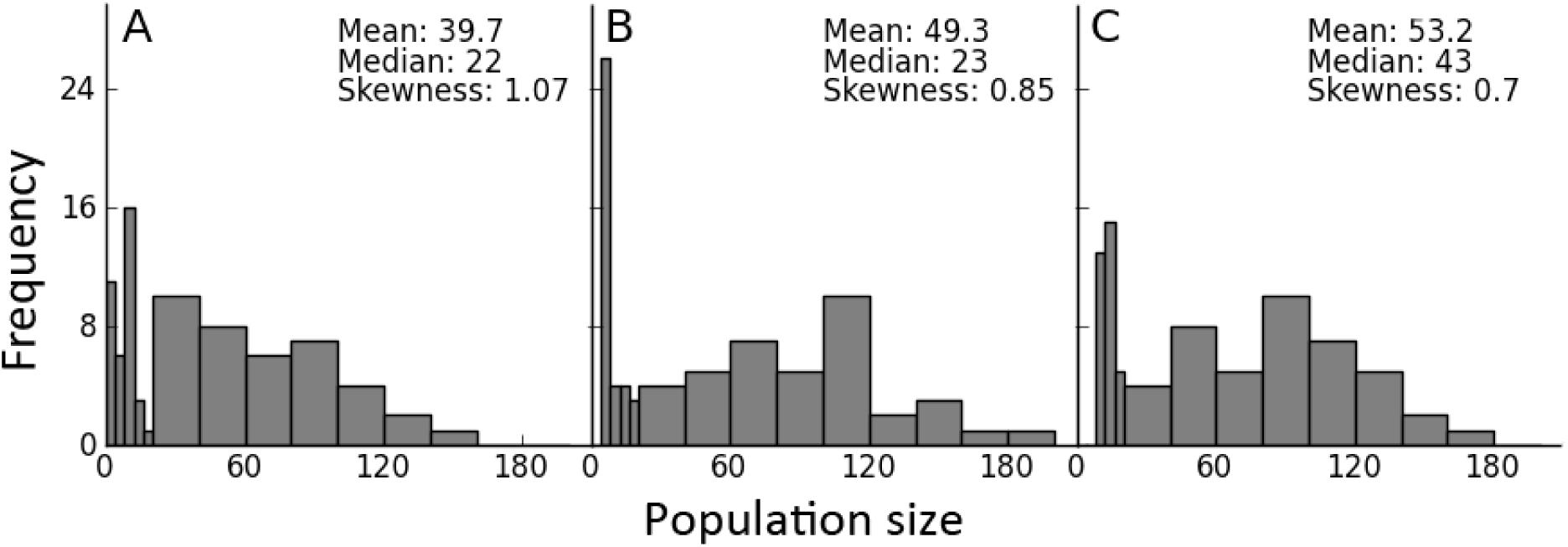
Population size distributions of the unperturbed and LLC-treated populations. A. Unperturbed (C2). **B.** L1 (lower threshold=4). **C**. L2 (lower threshold=10). When the population size is between 0-20, bin size is 4 and 20 otherwise. LLC treatments did not have a major effect on the population size distribution.

**Figure 5.**
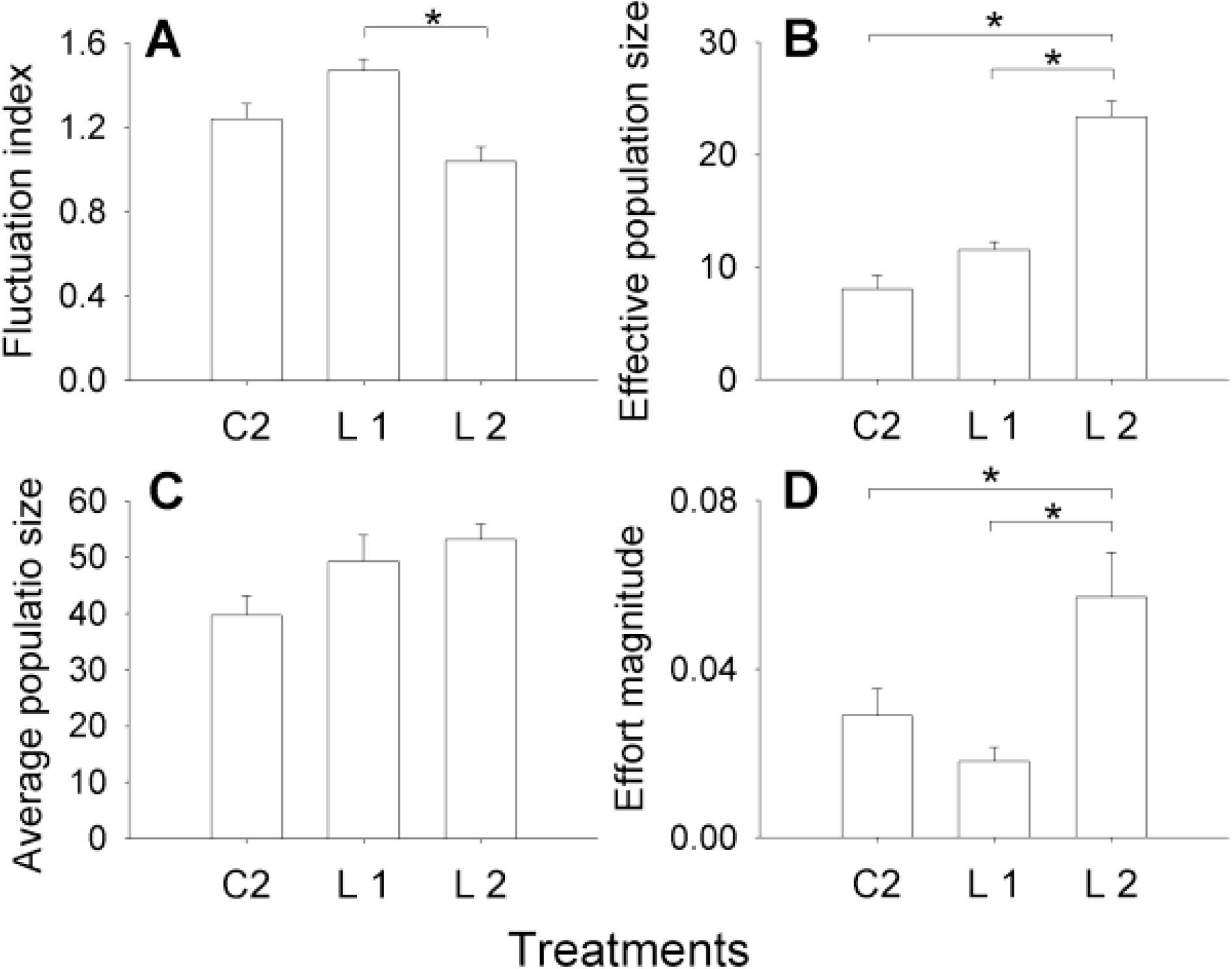
Dynamics after applying LLC. C2 represents the unperturbed populations while L1 and L2 stand for LLC thresholds of 4 and 10 respectively. Each bar represents a mean over 5 replicate populations. Error bars denote standard error around the mean and * denotes *p* < 0.05. **A. Fluctuation index**: Although there is a significant difference between L1 and L2, neither of them is significantly different from C2. L2 significantly increased **B**. **Effective population size** but none of the LLC treatments affect **C**. **Average population size** significantly. **D. Effort magnitude:** L2 required significantly more external perturbation than both C2 and L1.

## 4. DISCUSSION

This study compares how the upper and lower limiter control methods affect the dynamics of biological populations. However, it is not a direct comparison of the efficiency of the two methods. This is because simulations (Tung et al. 2014) indicate that almost every control method is capable of inducing almost any level of constancy stability, depending on the value of the control parameter (i.e. *U* or *L*). As it is not possible to equate over the parameter values for two methods as different as ULC and LLC, it becomes meaningless to compare their efficiencies. A recent simulation study circumvented this problem by numerically figuring out the magnitude of perturbation needed by each control method to achieve an arbitrary pre-determined level of stability (Tung et al. 2014). The efficiencies of the control methods were then investigated at those parameter values by comparing the resultant effective population size, effort magnitude, etc. Clearly, such an exercise is logistically very difficult in the context of an experimental study, where one is severely constrained in terms of how many treatments can be investigated. Therefore, in this study, we limit ourselves to mechanistic understanding of how each control method functions and refrain from a direct comparison of their efficiencies in attaining stability.

### 4.1 Upper Limiter Control (ULC)

#### Population size distribution and FI

ULC affects the dynamics of fluctuating *Drosophila* population in two ways. Firstly, by definition, the breeding population size was not allowed to go beyond a pre-determined threshold. Secondly, reduction in the number of breeding adults in generation *t* decreased the number of eggs laid in the next generation (i.e. *t+1*), which in turn lowered the larval competition. As a result, there was an increase in the larval survivorship which reduced the occurrences of population crashes in generation *t+1*. This is demonstrated by the fact that application of ULC (Fig 1B and 1C) reduced the frequency of the lower population sizes compared to the unperturbed case (Fig 1A). Consequently, the distribution of population sizes became more symmetric, the median approached the mean and the skewness reduced. This result is the biological analogue of the truncation of the stock-recruitment curve demonstrated by previous numerical studies (Figure 3.10 in Hilker and Westerhoff 2005) and provides a biological, mechanistic understanding of how ULC enhances the constancy stability of populations (Fig 2A). The reduction in population FI due to ULC is consistent with an earlier study on ciliates that had reported reduced variability in population size upon imposing an upper threshold (Fryxell et al. 2005).

#### Effective population size

We also investigated the potential impact of ULC on the genetic stability of populations by quantifying the effective population size (*N*_*e*_). *N*_*e*_ is defined as the corresponding size of an ideal population which loses heterozygosity at the same rate as the given population and is calculated here as the harmonic mean of the population sizes over time (Allendorf and Luikart 2007). Since the rate of loss of heterozygosity has an inverse relationship with *N*_*e*_, it follows that higher values of *N*_*e*_ are more desirable for enhanced genetic stability of populations. Also, as the harmonic mean gets affected more by the lower values in a time series, *N*_*e*_ is sensitive to population bottlenecks. Since ULC reduces population crashes, it was expected to enhance *N*_*e*_. However, although both levels of ULC increased *N*_*e*_ compared to the unperturbed treatment (Fig 2B), the difference was not statistically significant. Moreover, in spite of medium to large effect sizes of the differences between the means, the confidence intervals around the effect sizes were wide (Table S2). Together these suggest that there was wide variation in terms of the ability of ULC in enhancing *N*_*e*_. This is primarily because although ULC reduces the frequency of population crashes, it cannot ameliorate it completely (Fig 1B and 1C) and populations can still hit low values through demographic or environmental stochasticity. This observation is also consistent with a recent study which found that ULC was the least effective among six control methods in terms of increasing *N*_*e*_ (Tung et al. 2014). To summarize, ULC is not a reliable method in terms of enhancing *N*_*e*_.

#### Average population size

ULC also failed to affect the average population size of the controlled populations (fig 2C). Although removal of individuals is expected to reduce the population size, control methods that involve culling can sometimes lead to an increase in average population size. This phenomenon has been called the paradox of limiter control (Hilker and Westerhoff 2006) or the Hydra effect (Abrams 2009) in the ecological literature. It happens because culling reduces the negative effects of density dependence on realized growth rate, thus off-setting the loss of numbers due to removal. However, when the stock-recruitment curve of a population shows a long flat tail, as can be inferred for our populations from Fig 1A, then the hydra effect is not expected for a large range of ULC values. This is because for such populations, until and unless the ULC thresholds are fairly low, the increase in the realized growth rate is not sufficient to cause an increase in the mean population size after compensating for the number of adults removed. This phenomenon has been observed in earlier theoretical studies (Hilker and Westerhoff 2005) and was also captured in our simulations (Fig S3C).

#### Effort magnitude

Both U1 and U2 required effort to the tune of 40-50% of the corresponding average population size (Fig 2D). This is because although a fixed upper threshold restricts the number of breeding females, the high fecundity of the flies ensured a substantially higher-than-threshold population size in the next generation. Consequently, ULC had to be applied in most generations which translated into a high effort magnitude. This observation is consistent with a recent theoretical investigation which found that ULC entailed a relatively high effort magnitude (Tung et al. 2014). Our results show that when cost of culling is high, ULC is likely to be economically undesirable.

#### Persistence

Due to its tendency to diminish the frequency of population crashes, ULC reduced the extinction probability of the controlled populations, although the effect was statistically marginally insignificant (Fig 3A). It has already been suggested that the presence of some kind of upper threshold in population size is needed to optimize the yield and reduce the extinction probability of harvested populations in unpredictable environments (Lande et al. 1997). Our data demonstrates that ULC is also capable of enhancing the persistence of non-exploited populations.

To summarize, our results indicate that although ULC induces constancy and persistence stability, it is not very efficient in terms of enhancing the genetic stability of populations and is economically expensive.

### 4.2. Lower Limiter Control (LLC)

A numerical comparison of six control methods had concluded that LLC was the optimal control method under most (though not all) circumstances (Tung et al. 2014). Therefore, we investigated LLC under slightly more stringent conditions than ULC. Since under most real-life conditions, one is forced to operate on estimates of population sizes, rather than precise counts, we explicitly incorporated some degree of imprecision in our implementation of LLC. For this, rather than using the exact counts of female numbers in the population, we followed a previous protocol (Dey and Joshi 2006) and estimated the number of females under the assumption of 1:1 sex ratio. Since this assumption will evidently be not met in every generation, some amount of estimation error is built into the experimental procedure.

#### Population distribution and constancy stability

LLC directly prevents population sizes from going below a threshold without placing any restrictions on how high they can get. Consequently, although the population never hits the lower values in Fig 4B and 4C, there is not much change in the long right hand tail, and the skew values reduce more slowly with increase in the threshold. Compared to ULC (Fig 1B and 1C), the spread remains considerably wider. Not surprisingly therefore, the FI of neither L1 nor L2 were significantly different from the corresponding unperturbed population (Fig 5A). This failure to reduce FI could indicate that LLC does not work in real populations. It could also be attributed to the fact that we incorporated some degree of imprecision in the application of LLC. Either way, our results indicate that a lot more empirical studies are needed to establish the efficacy of LLC in stabilizing biological populations.

#### Effective and average population size

Although LLC treatments could not enhance the constancy stability, L2 increased the *N*_*e*_ significantly (fig 5B). This is important as low *N*_*e*_, and thereby enhanced chance of loss of genetic diversity, has been shown to have detrimental effect on population sustenance (Newman and Pilson 1997). However, this attainment of higher *N*_*e*_ is trivial as restocking methods increases the harmonic mean by ensuring that the population size never goes below a lower threshold. More crucially, in spite of restocking, there was no statistically significant effect of LLC on the average population size (Fig 5C) which contradicts earlier theoretical predictions (Hilker and Westerhoff 2005). Interestingly though, the effect sizes of all pair-wise differences were large (Table S3) and Fig 5C does reveal an increasing trend in average population size with increase in intensity of LLC. The lack of statistical significance is better explained by the presence of large amount of variation among replicates in the L1 and L2 treatment, which might be an artefact of the noise incorporated during census. Thus the effect of LLC on average population size merits further investigation.

#### Effort magnitude

The effort magnitude required for LLC is much, much less compared to that of ULC (*cf* Fig 2D and 5D and note the difference in scale). Unfortunately, the two values are not directly comparable here, since they lead to different amounts of constancy stability. However, an earlier comparison of control methods have shown that for attaining comparable levels of stability, LLC typically requires the least amount of effort (Tung et al. 2014). This leads to the possibility that the performance of LLC can be probably improved significantly without incurring too much cost in terms of effort magnitude.

#### Persistence

Although LLC failed to enhance constancy, it had a significant effect in terms of persistence (Fig 3B) which was almost comparable to that of ULC (*cf* 3A and 3B). *Prima facie*, this is counterintuitive as LLC does not involve culling and hence cannot reduce the frequency of population crashes to zero or very low numbers. This in turn means that it is not expected to reduce extinction probability which was scored before the imposition of perturbation. The discrepancy gets resolved when we recall that there are two ways in which a population can go extinct. The first is when the numbers in generation *t* crash to zero due to overpopulation in generation *t-1*. The second is when overpopulation in generation *t-1* causes the numbers to become very low in generation *t*, which in turn leads to extinction in generation *t+1*. While LLC cannot ameliorate the first kind of extinction, it can reduce the probability of the second kind due to restocking by fecund individuals. This is the converse of ULC, which can only reduce the first kind of extinction but not the latter type. This leads to the prediction that the optimal method to promote persistence would depend on the kind of extinction suffered by the target population. For any population which, like our *Drosophila* cultures, suffer from both kinds of extinctions, a control method that incorporates both culling and restocking would be the most effective in terms of promoting persistence.

To summarize, the present experiments are inconclusive about the effects of LLC on constancy. However, it is clear that LLC boosts effective population size and persistence while requiring relatively less effort.

### 4.3 Caveats

Several caveats need to be considered while extrapolating our results to natural conditions. In our study, we consider culling and restocking to be equivalent in terms of effort magnitude. However, under some circumstances, killing individuals in a population might be less expensive than reintroduction from a source and vice versa. In all those cases, the effort magnitude needs to be suitably scaled to arrive at the actual economic cost. Another fact not considered here is that the economic cost of census also might be very different for ULC and LLC. In the former case, the entire population needs to be counted while in the latter case, census effort is restricted to the point that the threshold number of organisms is cited. Furthermore, although we have calculated effective population size solely as a function of the census size, in reality, culling and restocking differ in how they affect the loss of genetic variation in a population. This is because if the individuals used in restocking belong to a different gene pool, then there would be relatively lower loss of genetic variation than what is expected based on the values of effective population size. Finally, we have only examined the effects of ULC and LLC on spatially-unstructured populations of a given species, whereas most natural populations exist in spatially-structured communities of many species. Thus, several factors need to be taken into account before our experimental results can become usable in real-life scenarios. However, given that erroneous application of control methods can lead to ecological disasters (Pyke 2008), studies such as ours can help in bridging the gap between theory and practical applications.

## 5. CONCLUSIONS

Our empirical results suggest that ULC is an efficient method in terms of reducing fluctuations as well as extinction probability. However, it does not increase the effective population size and needs fairly large magnitude perturbations. On the other hand, although the efficacy of LLC in reducing population fluctuations is equivocal, it increases effective population size, reduces extinction probability and requires relatively less effort. Thus, the choice of methods under a given condition would depend upon the aspect of the dynamics that needs to be stabilized. We also investigated the generalizability of our empirical results. Our Ricker-based simulations did not contain any species-specific parameters, and yet were able to corroborate most of the empirical observations. This indicates that our experimental results are not due to some specific features of *Drosophila* biology, and hence, likely to be broadly applicable. It has been theoretically shown that populations that experience scramble competition and are randomly distributed over space, follow the Ricker dynamics (Brännström and Sumpter 2005). Since large number of organisms belonging to diverse taxa fulfil these two conditions, our results are likely to be relevant in wide-ranging scenarios. However, there are several caveats to our results, and any extrapolation to real-life scenarios must be supplemented by relevant system-specific information.

## ACKNOWLEDGEMENTS

We thank Joseph Paul Salve and Dhanashri Nevgi for help in running the experiments. ST thanks Council for Scientific and Industrial Research, Government of India for financial support through a Senior Research Fellowship. AM thanks Department of Science and Technology, Government of India for financial support through a KVPY fellowship. This study was supported by a research grant from the Council for Scientific and Industrial Research, Government of India and internal funding from IISER-Pune.

## Supplementary material

### S1: Description of simulations

The dynamics of the populations were simulated using the Ricker model (Ricker 1954). This model is given as *N*_*t*+1_=*N*_*t*_*\*exp(r*(1-N_t_/K))*, where *r, K* and *N*_*t*_ denote intrinsic growth rate, carrying capacity and population size at time *t* respectively. Upper and lower limiter control were imposed according to the following mathematical form (Tung et al. 2014):

Upper Limiter Control (ULC): *a*_*t*_ = *min [b_t_, H]*
Lower Limiter Control (LLC): *a*_*t*_ = *max [b_t_, h]*

where *b*_*t*_ and *a*_*t*_ are the population sizes before and after perturbation in the *t*^*th*^ generation. *b_t+1_ = f(a_t_)*, where *f* stands for Ricker model. *H* and *h* are upper and lower threshold of ULC and LLC respectively.

#### Parameter values

We fit the Ricker model to the unperturbed experimental time-series (C1 and C2) and obtained a mean *r* and *K* of 2.6 and 34 respectively. These values of *r* and *K* were then used to simulate the basic dynamics of unperturbed and controlled populations. Matching the experimental values, upper threshold for U1 and U2 were kept at 15 and 10, whereas lower threshold for L1 and L2 were 4 and 10 respectively.

#### Stochasticity and lattice effect

Noise can significantly influence the dynamics of perturbed populations (Dey and Joshi 2007). Therefore we incorporated noise to both the parameters, *r* and *K*, in each iteration. For this, we picked the *r* and *K* values from uniform distributions of 2.6±0.5 and 34±15 respectively. Real organisms always come in integer numbers, and incorporating this in simulations can affect the dynamics of the systems; a phenomenon termed as lattice effect (Henson et al. 2001, Domokos and Scheuring 2004). We accounted for lattice effect by rounding off the number of organisms in each generation, as well as the values of the perturbations, to the nearest integer.

The magnitudes of perturbations were computed for each of the treatments of the control methods after assuming a 1:1 sex ratio. Thus, for example, when the lower limit of LLC was set to 4, the control method was implemented only when the population size fell below 8. Although the Ricker model does not take zero-values and hence, can never show extinction, the same is not true for the integerized model. Therefore, we set reset rules similar to the experiment for our simulations. When the unperturbed populations and ULC treatments went extinct, the population size was reset to 8. No resets were necessary for the LLC treatments where the control method automatically ensured reset. Each simulation was run for 100 iterations and FI, average population size and effective population size were computed over the resulting time series. All figures (Fig S5 and S7) represent means over 100 independent replicates for each treatment.

**Table S2.** **Tukey p value and effect size for all pairwise combinations among C1, U1 and U2**

| ULC | Pairwise combinations | Tukey p | Effect size ± 95% CI |
| --- | --- | --- | --- |
| Fluctuation index | C1-U1 | 0.0007 | $3.66 \pm 2.03$ |
| | C1-U2 | 0.0004 | $3.38 \pm 1.93$ |
| | U1-U2 | 0.78 | $0.43 \pm 1.25$ |
| Av. population size | C1-U1 | - | $0.95 \pm 1.31$ |
| | C1-U2 | - | $0.85 \pm 1.29$ |
| | U1-U2 | - | $0.17 \pm 1.24$ |
| Effective population size | C1-U1 | - | $1.17 \pm 1.34$ |
| | C1-U2 | - | $1.49 \pm 1.4$ |
| | U1-U2 | - | $0.23 \pm 1.24$ |
| Effort magnitude | C1-U1 | 0.0002 | $5.39 \pm 2.67$ |
| | C1-U2 | 0.0002 | $6.00 \pm 2.91$ |
| | U1-U2 | 0.08 | $1.25 \pm 1.35$ |
| Extinction probability | C1-U1 | 0.21 | $1.4 \pm 1.38$ |
| | C1-U2 | 0.05 | $1.8 \pm 1.47$ |
| | U1-U2 | 0.7 | $0.48 \pm 1.26$ |

**Table S3.** **Tukey p value and effect size for all pairwise combinations among C2, L1 and L2**

| LLC | Pairwise combinations | Tukey p | Effect size ± 95% CI |
| --- | --- | --- | --- |
| Fluctuation index | C2-L1 | 0.11 | $1.43 \pm 1.39$ |
| | C2-L2 | 0.17 | $1.14 \pm 1.34$ |
| | L1-L2 | 0.004 | $2.85 \pm 1.76$ |
| Av. population size | C2-L1 | - | $0.93 \pm 1.31$ |
| | C2-L2 | - | $1.71 \pm 1.45$ |
| | L1-L2 | - | $0.41 \pm 1.25$ |
| Effective population size | C2-L1 | 0.17 | $1.43 \pm 1.39$ |
| | C2-L2 | 0.0002 | $4.7 \pm 2.4$ |
| | L1-L2 | 0.0003 | $4.26 \pm 2.24$ |
| Effort magnitude | C2-L1 | 0.58 | $0.93 \pm 1.31$ |
| | C2-L2 | 0.05 | $1.44 \pm 1.39$ |
| | L1-L2 | 0.008 | $2.21 \pm 1.57$ |
| Extinction probability | C2-L1 | 0.06 | $2.4 \pm 1.63$ |
| | C2-L2 | 0.02 | $2.8 \pm 1.74$ |
| | L1-L2 | 0.78 | $0.37 \pm 1.25$ |

**Figure S4.**
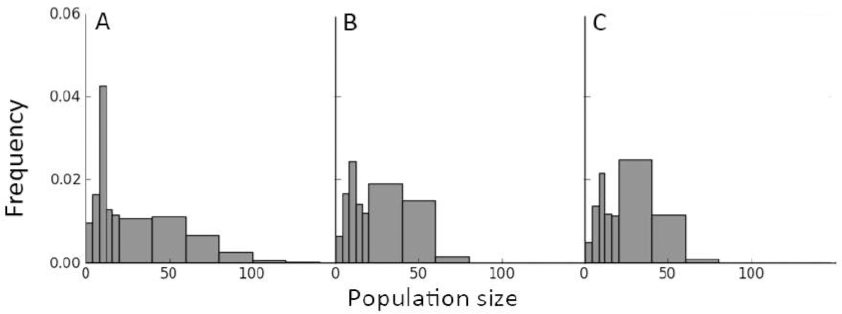
Size distributions of the simulated unperturbed and ULC-treated populations. **A.** Unperturbed (C1). **B.** U1 (upper threshold=15). **C**. U2 (upper threshold=10). When the population size is between 0-20, bin size is 4 and 20 otherwise.

**Figure S5.**
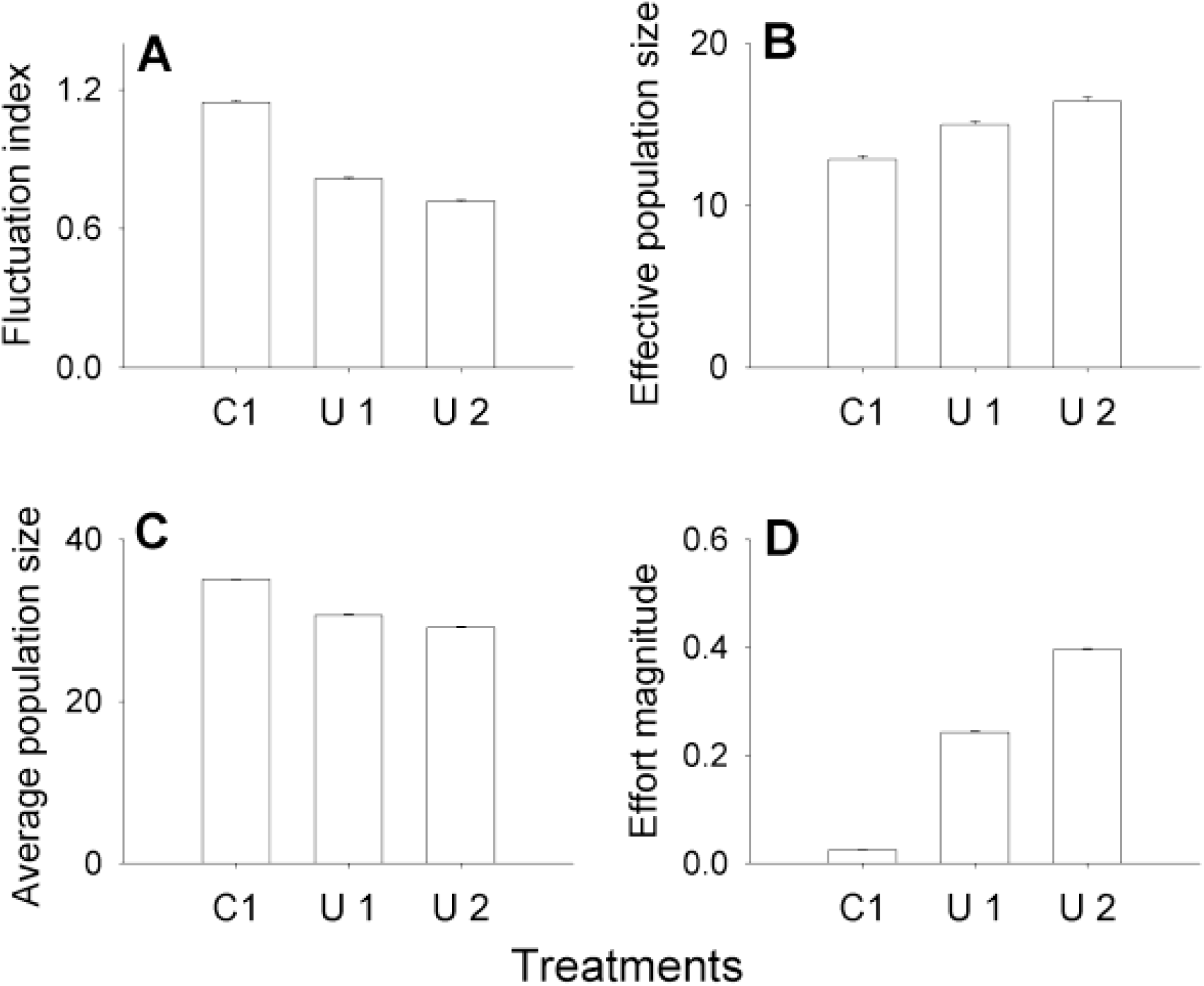
Simulated dynamics after applying ULC. C1 represents the unperturbed populations while U1 and U2 stand for ULC thresholds of 15 and 10 respectively. Each bar represents a mean over 5 replicate populations. Error bars denote standard error around the mean. **A. Fluctuation index**, **B. Effective population size**, **C. Average population size and D. Effort magnitude.**

**Figure S6.**
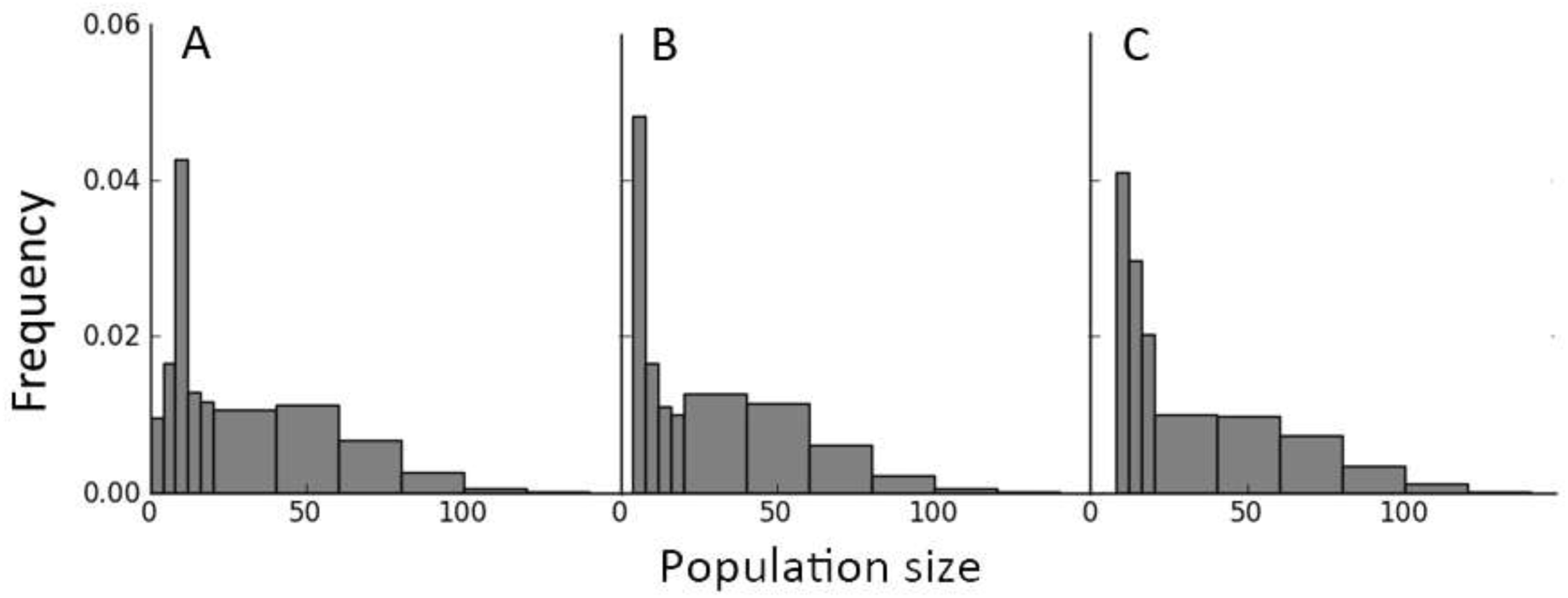
Size distributions of the simulated unperturbed and LLC-treated populations. A. Unperturbed (C2). **B.** L1 (lower threshold=4). **C**. L2 (lower threshold=10). When the population size is between 0-20, bin size is 4 and 20 otherwise.

**Figure S7.**
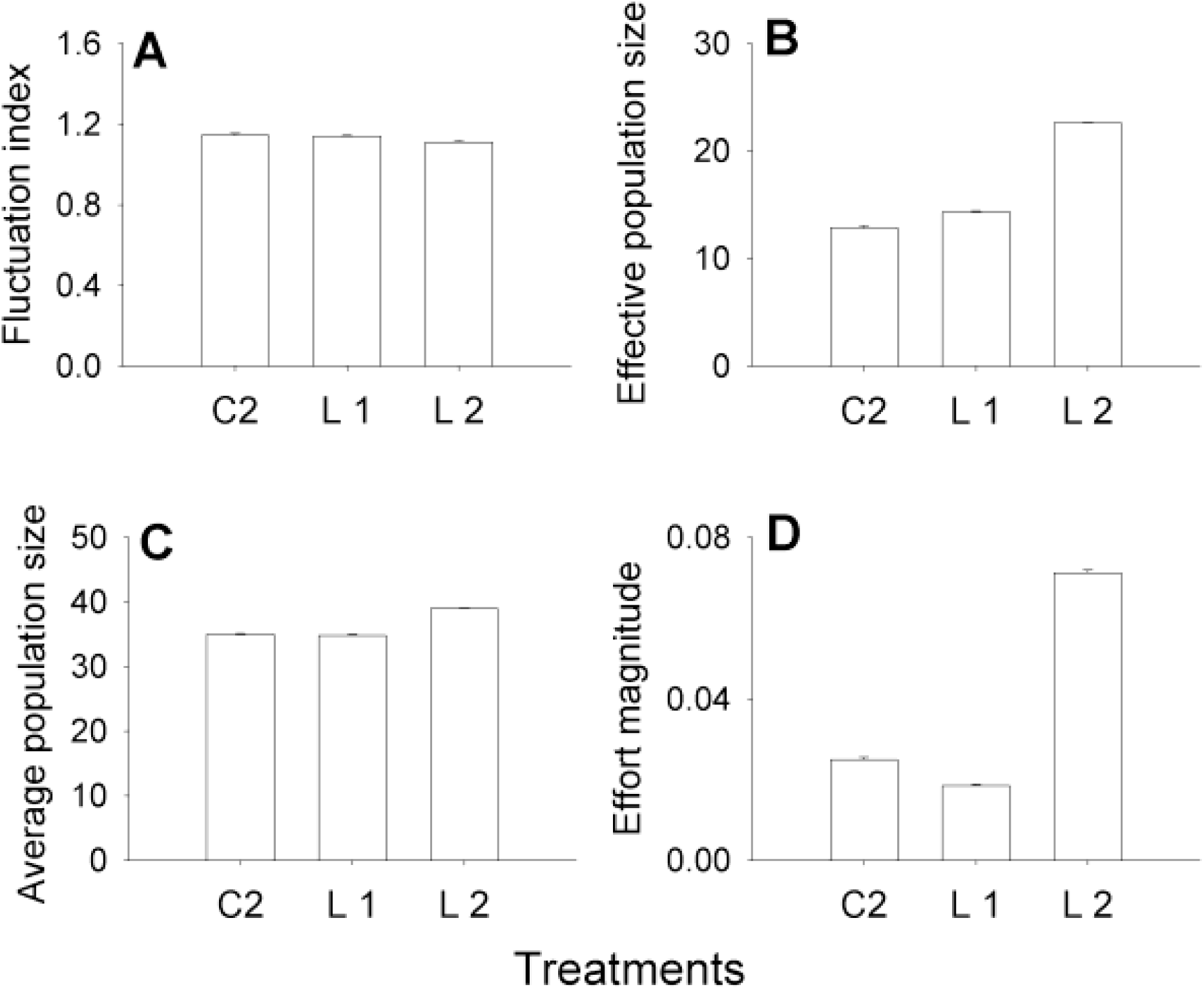
Simulated dynamics after applying LLC. C2 represents the unperturbed populations while L1 and L2 stand for LLC thresholds of 4 and 10 respectively. Each bar represents a mean over 5 replicate populations. Error bars denote standard error around the mean. **A. Fluctuation index**, **B. Effective population size**, **C. Average population size and D. Effort magnitude.**

